# Correlating surface plasmon resonance microscopy of living and fixated cells with electron microscopy allows for investigation of potential preparation artifacts

**DOI:** 10.1101/817288

**Authors:** Eva Kreysing, Silke Seyock, Hossein Hassani, Elke Brauweiler-Reuters, Elmar Neumann, Andreas Offenhäusser

**Affiliations:** Forschungszentrum Jülich GmbH, Institute of Complex Systems, Bioelectronics (ICS-8), 52425 Jülich, Germany; University of Cambridge, Department of Physiology, Development and Neuroscience (PDN) Downing Street, CB2 3DY Cambridge, UK; Klinik und Poliklinik für Anästhesiologie und Intensivmedizin, AG IRIS, Universitätsmedizin Rostock Schillingallee 69, 18057 Rostock, Germany; Forschungszentrum Jülich GmbH, Helmholz Nanofacility (HNF), 52425 Jülich, Germany

**Keywords:** Cell-substrate distance, Surface Plasmon Resonance Microscopy, Electron Microscopy, Fixation artifacts

## Abstract

The investigation of the cell-substrate interface is of great importance for a broad spectrum of areas such as biomedical engineering, brain-chip interfacing and fundamental research. Due to its unique resolution and the prevalence of instruments, electron microscopy (EM) is used as one of the standard techniques for the analysis of the cell-substrate interface. However, possible artifacts that might be introduced by the required sample preparation have been the subject of speculation for decades. Due to recent advances in Surface plasmon resonance microscopy (SPRM), the technique now offers a label-free alternative for the interface characterization with nanometer resolution in axial direction. In contrast to EM, SPRM studies do not require fixation and can therefore be performed on living cells. Here, we present a workflow that allows us to quantify the impact of chemical fixation on the cell-substrate interface. These measurements confirmed that chemical fixation preserved the average cell-substrate distances in the majority of studied cells. Furthermore, we were able to correlate the SPRM measurements with EM images of the cell-substrate interface of the exact same cells allowing us to identify regions with good agreement between the two methods and reveal artifacts introduced during further sample preparation.

## 1 Introduction

Understanding the interaction of cells with artificial materials is essential for fundamental research as well as biomedical applications such as cell and tissue engineering, the design of implants and prosthesis but also in the development of brain-computer interfaces. [1, 2, 3, 4, 5] Surface modifications such as nanostructures and coatings can act as chemical, mechanical and topological cues that influence cell-substrate adhesion, cell migration and even stem cell fate. [6, 7, 2] In the development of implants such as retina implants, a strong cell-substrate adhesion is crucial for a good cell-electrode communication while mediating a good integrity. [8, 9, 10] Understanding said interactions requires sophisticated imaging techniques with a resolution in the nanometer range. This makes electron microscopy (EM) an indispensable tool in the field. EM includes multiple different techniques such as transmission electron microscopy (TEM) [11, 12, 13, 14] and scanning (transmission) electron microscopy (S(T)EM). [15, 16, 17, 18, 19, 20, 21, 22, 23, 24, 25, 26, 27, 28] While S(T)EM can reach nanometer resolution, it requires extensive sample preparation prior to the FIB sectioning. As a first step, the cells are usually fixated using a cross linker such as glutaraldehyde which crosslinks proteins and preserves the cell structure. Volumetric studies showed that depending on the protocol, glutaraldehyde fixation might either preserve or change the cell volume in single cells [29] whereas shrinkage was reported when fixating corneal endothelial tissue. [30] However, the impact on the cell-substrate distance in the nanometer range has not been studied. Two frequently used preparation techniques are critical point drying (CPD) and resin embedding [31, 32, 21, 7]. In CPD, the chemical fixation is followed by a dehydration of the cell. Intracellular water is exchanged with ethanol and subsequently replaced with liquid CO_2_, which is afterwards transferred into a supercritical fluid and blown off. This can be followed by heavy metal stainings which allow for resolution of the cellular ultrastructure in EM.

In resin embedding protocols, chemical fixation is often followed by heavy metal stainings e.g. with OsO_4_ and uranyl acetate. Subsequently, the intracellular water is replaced by a solvent e.g. ethanol and afterwards by resin which mechanically stabilizes the cell structure during FIB sectioning. In order to identify individual cells in EM, excess resin needs to be removed, preserving only the resin within the cell and a thin layer on top which is afterwards polymerized by heat (thin layer plastification (TLP)). Whilst CPD can be carried out relatively quickly, resin embedding is usually more time-consuming and often requires the handling of toxic and radioactive substances. The two methods also differ regarding the artifacts that they introduce. On the one hand, CPD can produce drying artifacts like volume shrinkage of the cells [33, 34, 35, 36] and a porous structure at the interface. [21, 23, 37] On the other hand, the exchange of water against a solvent in resin embedding protocols may produce dehydration artifacts which might be less obvious in EM images because the resin only stabilizes the structure that has been obtained after the dehydration step. [38, 21]

There are several optical techniques, that in contrast to EM offer the opportunity to investigate the interface of the living cell. Examples are fluorescence interference contrast microscopy (FLIC) [39, 40] and metal induced energy transfer (MIET). [41] These techniques can be used to study the interface between the fluorescent labeled cell membrane and the specific substrate. The calculation of the cell-substrate distance in FLIC and MIET is based on the modulation of the fluorescence intensity and the fluorescence lifetime of the dye molecules in the vicinity of specific substrates, respectively.

While the axial resolution can reach nanometer accuracy, the lateral resolution of FLIC and MIET has been reported to reach an accuracy of up to 900 *nm* and 200 *nm*, respectively. These techniques allow for insights into the living cell, but they might affect the cellular behavior as fluorescent dyes can be cytotoxic. [42, 43]

Surface plasmon resonance microscopy (SPRM) represents a label-free technique allowing for the study of the cell-substrate interface. Widefield SPRM is based on the illumination of the interface at a single but adjustable incidence angle, which enables imaging of the interface in real time. [44, 45, 46, 47, 48, 49, 50, 51, 52, 53, 54, 55, 56]. This has been demonstrated to resolve cell adhesion complexes as well as other subcellular structures. [50, 52, 57] In scanning SPRM, the incident light is focused at the cell-substrate interface, resulting in a spotwise sample illumination with a wide range of angles. Afterwards, the intensity of the reflected light over the angle spectrum is analyzed to quantitatively determine the cell-substrate distance as well as the intracellular refractive index (RI). Since scanning SPRM permits a label-free measurement of the cell-substrate distance in living cells with similar lateral resolution to MIET while avoiding phototoxicity, it has been proven to be a valuable alternative to the above optical techniques [44, 58, 49]. Due to recent improvements to the analysis routine, SPRM can now also reach an accuracy of up to 1.5 *nm* in axial direction, [56] which is comparable to FLIC, MIET and EM.

By performing a fixation protocol on a set of cells and measuring each individual cell with SPRM both prior to and after fixation as well as with FIB-S(T)EM, we were able to measure the impact of the chemical fixation on the cell-substrate interface of the investigated cells and to identify artifact affected regions. Such a combination of SPRM and FIB-S(T)EM holds a huge potential, since SPRM can characterize the unperturbed living cell and may even give insights into cellular dynamics [49, 59, 53, 52, 54, 56] while EM images of the FIB-cross sections allow us to resolve the cell membrane at focal adhesion points of the fixed cell with nanometer resolution in x- and z-direction.

## 2 Results

For this study, we used an objective-based SPRM setup which provides a live-imaging as well as a scanning mode for observing cells cultured on gold-coated glass samples. The live-imaging mode (see Figure 1a) is a widefield imaging technique which allows us to observe the cell-substrate contact area qualitatively. Here, areas with a close cell-substrate distance appear relatively bright whereas areas with higher cell-substrate distances and the surrounding of the cell appear darker (see Figures 1b, 2b, 2e). The scanning mode (see Figure 1c) is based on a point-wise illumination of the substrate and allows us to quantify the cell-substrate distance as well as the intracellular refractive index (RI) at each point. As described in greater detail in the Experimental Section, we capture one image of the back focal plane (BFP) at each scanning point, extract a reflectance profile (see Figure 1f) from this image and fit a layer model describing the sample structure (Figure 1 d). As a result, we can generate a 3D representations of the basal cell membrane as well as RI profiles of the cytosol [56].

**Figure 1:**
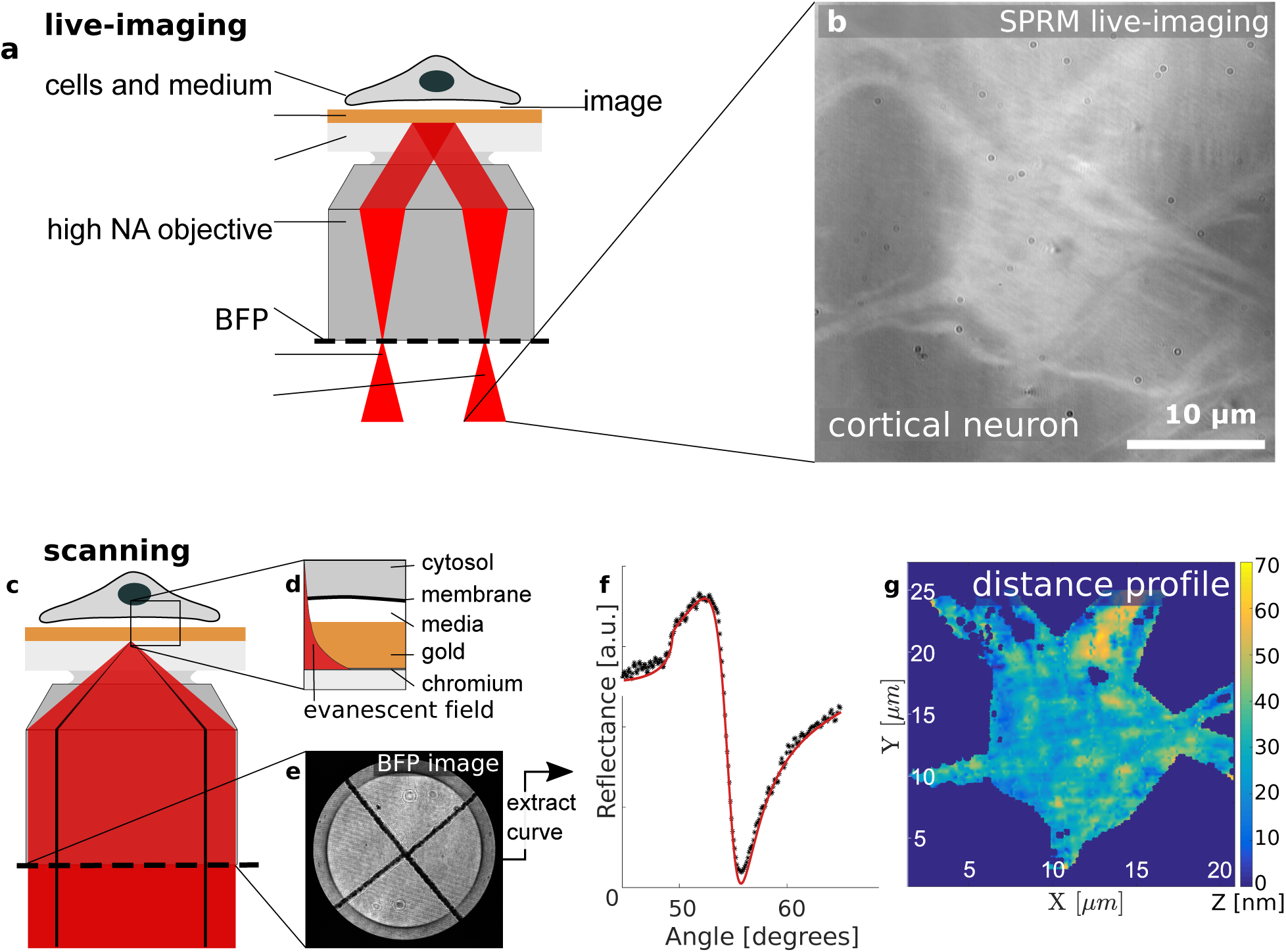
Qualitative and quantitative interface characterization. The SPRM live-imaging and scanning mode allow for a qualitative and a quantitative characterization of the cell-substrate interface, respectively. a) In the live-imaging mode, light is focused on the back focal plane (BFP) of the TIRF objective resulting in an illumination of the cell-substrate interface with parallel light. b) If illuminated under the resonance angle of the bare cell-substrate interface, areas of close cell-substrate distances appear relatively bright while areas with poor adhesion appear relatively dark. c) in the scanning mode, the entire BFP is illuminated with radially polarized light resulting in an illumination of the sample with a broad range of angles. d) Illuminating the sample under angles larger than the critical angle creates an evanescent field penetrating the cytosol. e) Recording the BFP at each scanning point allows for the extraction of the reflectance curve (f). f) Modeling the sample as a multilayer system, we can simulate the reflectance curve. Fitting the model to the data, we can extract the cell-substrate distance. g) Combining the data from the entire set of scanning points, we can reconstruct the 3D structure of the basal cell membrane.

**Figure 2:**
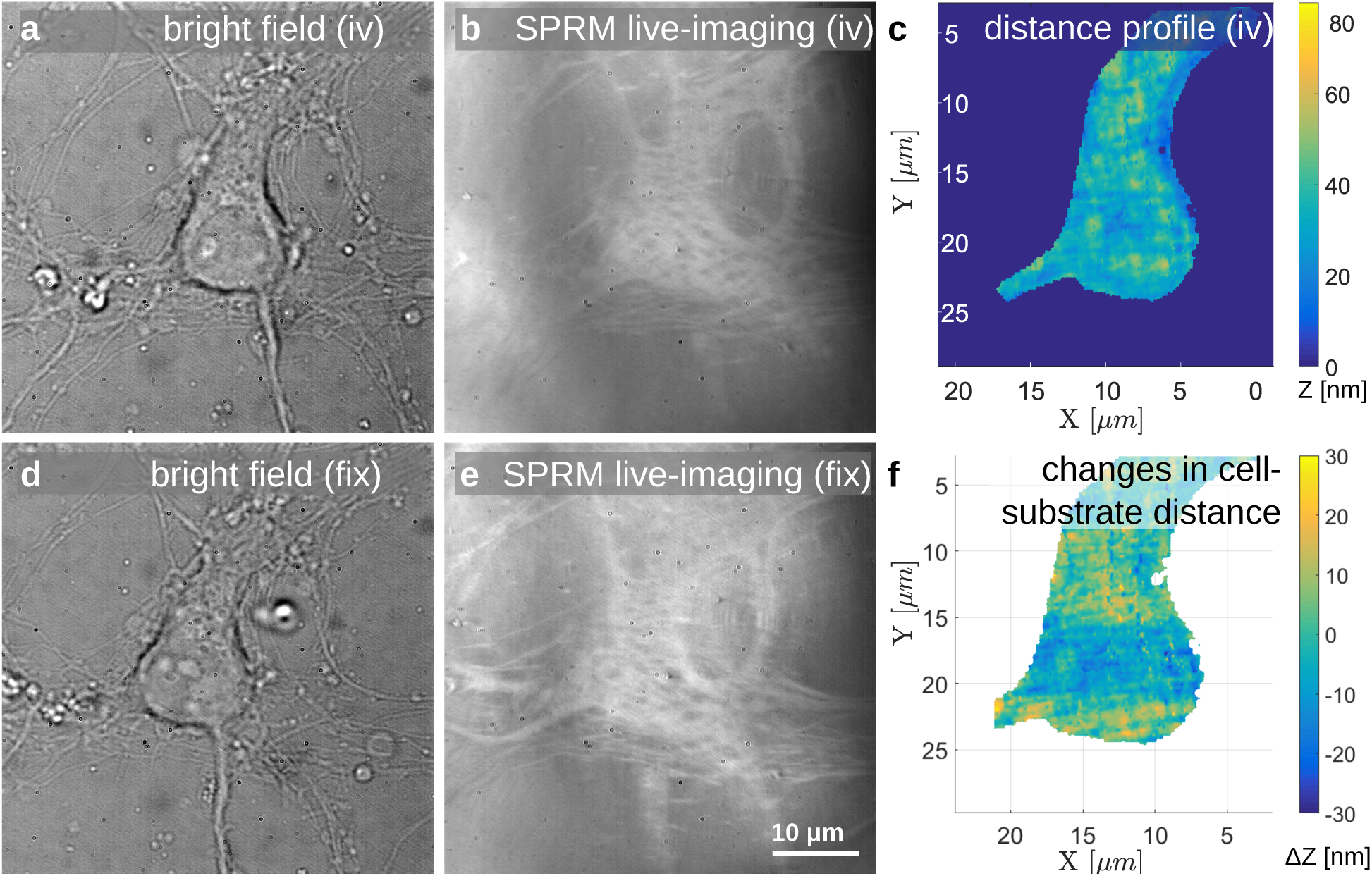
Fixation-induced changes at the interface observed with SPRM. a, d) Bright field images of the neuron give information regarding the shape of the soma and the dendrites *in vitro* (iv) and fixed (fix), respectively. b, d) The differences in contrast between the background and the adherent areas in the live-imaging of the living and the fixed cell indicate a change at the cell-substrate interface. c) The scanning results reveal the cell-substrate distances (results shown here correspond to the living cell). f) Subtracting the distance profiles of the living and the fixed system reveals the changes in the cell-substrate distances which are induced by the chemical fixation.

The cells used were primary cortical rat neurons cultured on poly-L-lysine (PLL) coated SPRM substrates and investigated on day in vitro (DIV) 7. We used the SPRM live-imaging mode to localize cells with a good contrast compared to their surrounding and identified their position on the chip using the photo resist markers (see Experimental Section) before scanning the respective living cells. Afterwards, we fixed the cells using glutaraldehyde, washed the samples carefully with phosphate-buffered saline (PBS) and remounted the samples on the setup. Subsequently, the previously scanned cells were localized using the live-imaging mode and scanned with identical settings. Afterwards, we stained the samples with osmium tetroxide (OsO4), which binds to the phospholipids in the membrane. Tannic acid is then used to improve the fixation and mediate binding of uranyl acetate which binds to proteins and lipids in the final staining. These steps stabilize the cell membrane and the high atomic weight of the heavy metals leads to a good contrast between the cell membrane and the surrounding resin in EM. [60] This was followed by resin infiltration, removal of the extracellular resin and a polymerization (see Experimental Section).

Using a FIB-SEM system, we localized the same cells previously characterized with SPRM using photo resist markers and deposited additional platinum layers on the region of interest (see Experimental Section). The FIB sectioning was started from one side and stopped when the center of the cell was reached. Afterwards, we imaged the cross section with the integrated SEM system and continued FIB sectioning from the opposite side in order to prepare a thin lamella from the center of the cell. Finally, we soldered said lamella to a lift-out-grid which was then attached to a STEM holder. This sample was mounted in an STEM (STEM-in-SEM system) which allowed us to image the cell-substrate interface with a higher resolution compared to the SEM images.

To determine the changes in cell-substrate distance due to chemical fixation, each SPRM scan was analyzed as described in the Experimental Section. The resulting distance profiles of the living and fixed cells were subtracted. For this task, additional software was written to allow us to align the areas scanned before and after fixation correcting for displacement and rotation of the sample caused by the second sample mounting (*align and compare* software see Experimental Section). The changes in the distance profiles as well as the RI profile can be displayed in multiple ways:

The area plots shown in Figure 2f and Figure 6f provide information regarding the fixation process in the individual cells. Areas with small variations in the cell-substrate distance indicate that the fixation process has preserved the original structure of the cell-substrate interface (see central areas of the cell in Figure 2f). Areas with large distance variations can indicate that the cell has partially detached from the substrate or collapsed locally during the process. Figure 6f shows strong changes in the cell-substrate distance of up to 60 *nm*, indicating that the fixation process did not conserve the original structure in the respective areas.

**Figure 3:**
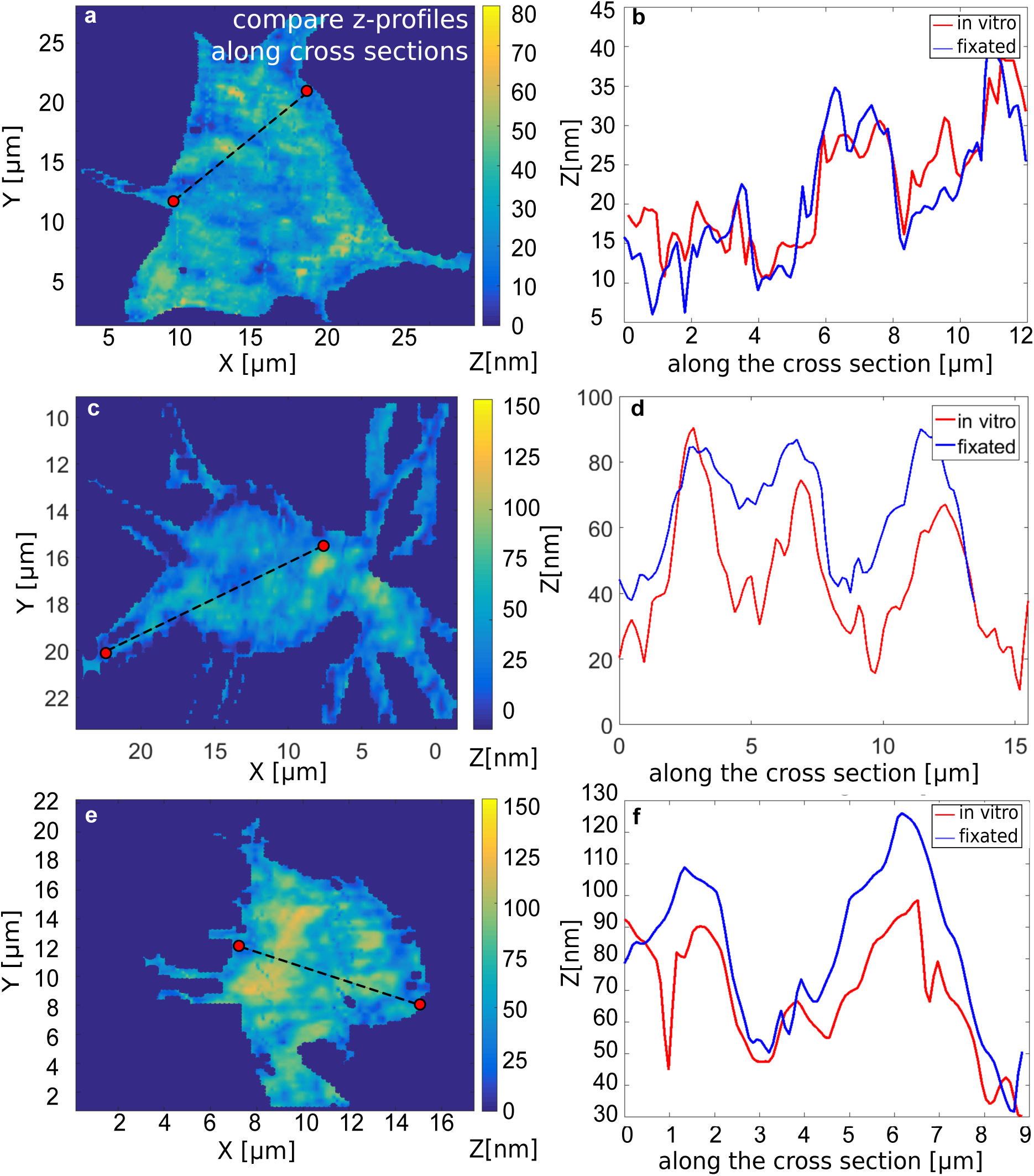
Distance profile along the cross section measured with SPRM. a, c, e) Definition of the cross-section via graphical input using the software *align and compare*. b, d, e) The cell-substrate distance is plotted along the chosen cross section in the states before and after the chemical fixation.

**Figure 4:**
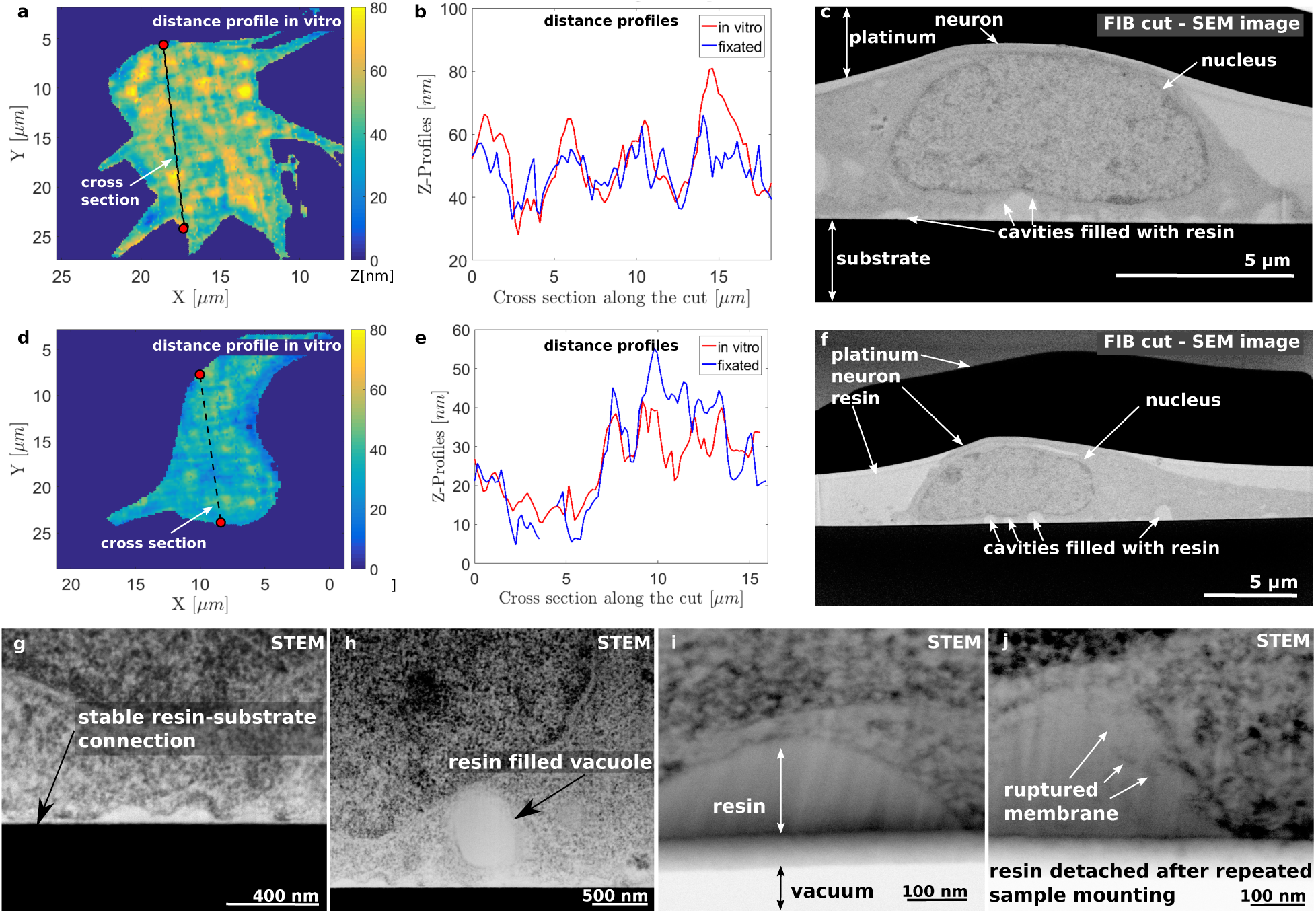
Correlation of SPRM and EM for two individual cells. The cell substrate distances are evaluated along the cross section with SPRM and FIB-SEM and STEM for the same cells and in the same direction. a, d) The cell-substrate distance is plotted as area scans representing the situation *in vitro*. The cross section is marked for the line-plot evaluations shown in (b, e). b,e) The distance is evaluated along the cross section marked in (a, d, respectively) for the system before and after the fixation. c, f) An SEM-image taken after FIB sectioning along the cross section marked in (a, d) reveals resin filled cavities at the cell-substrate interface. g) After extracting a lamella via FIB sectioning from the center of the cell shown in (d), the lamella was imaged with an STEM allowing for the resolution of the cell membrane. The close contacts found here are in good agreement with the SPRM results shown in (d, e) h) The high resolution of STEM allows us to discriminate between cavities at the interface and resin filled vacuoles as we see here. i) The STEM shows a cavity at the interface with an intact cell membrane. The resin was in contact with the substrate during the first mounting of the lamella (compare f). After the second mounting of the fragile lamella, the resin locally detached from the substrate. Therefore, the area underneath the resin appears white (vacuum) in the STEM (i, j). j) This cavity shows a ruptured cell membrane. The cell membrane might have been torn during the preparation process.

**Figure 5:**
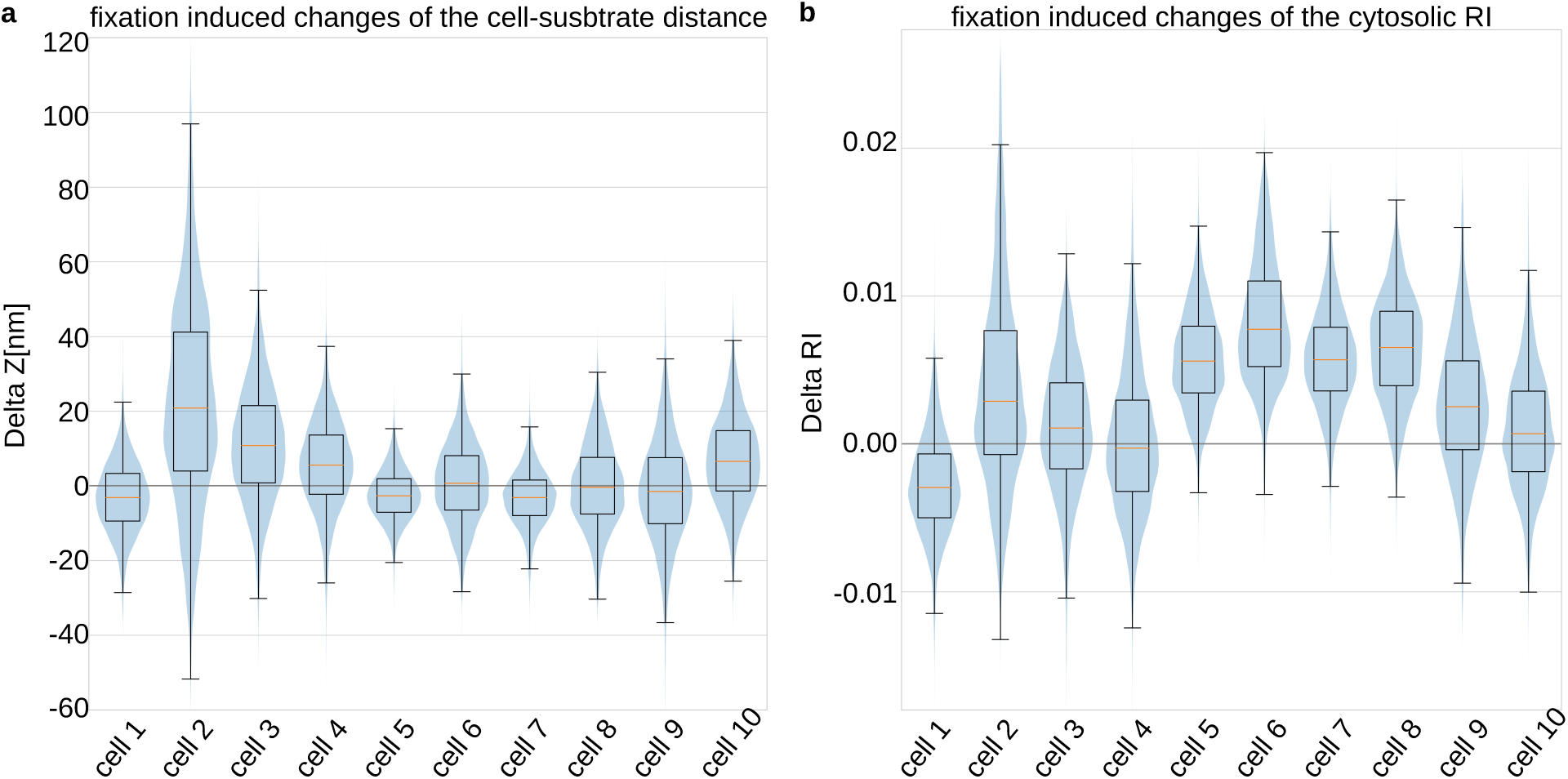
Fixation-induced changes in distance and refractive index. The changes in the cell-substrate distance and the cytosolic RI are evaluated for scanning point for 10 cells. a) We find prominent changes in the cell-substrate distance of cell 2 and cell 3 towards higher values while the remaining cells show random shifts. b) The cytosolic RI shows a systematic shift towards higher values after the fixation.

**Figure 6:**
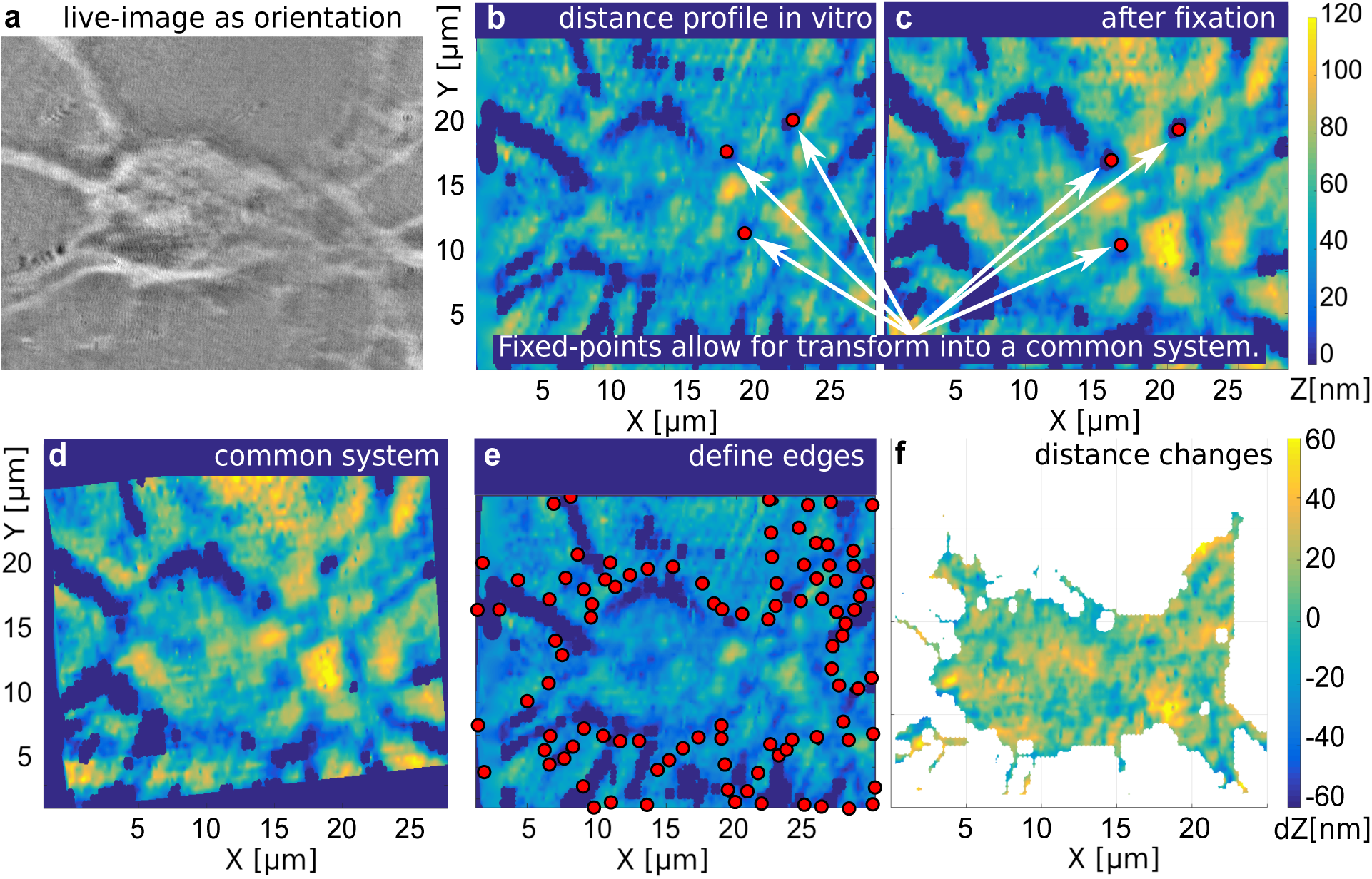
Working principle of the software *align and compare*. The software allows us to determine the fixation induced changes in the cell-substrate distance. a) A SPRM live-image of the cell allows us to identify the cell-adhesion areas. b, c) First, distance profiles before and after the fixation are shown in parallel and the user is asked to identify at least three salient points. d) Based on the coordinates of these points, a common coordinate system is determined. e) The edges of the cell adhesion area are defined by the user based on a brightfield image (a). f) The difference in the cell substrate distance before and after the fixation is determined and plotted as an area plot.

The line plots give quantitative information about the cell-substrate distance along a cross section between any two points. They enable us to compare the morphology of the cell in cross-section before and after fixation at a glance (Figure 3b, d, f). This facilitates the identification of fixation-related artifacts along the selected cross-section. These diagrams, which are chosen in the same direction as the cross-section in FIB sectioning, provide good comparability with EM images of the cell-substrate interface (Figure 4b, e). This in turn makes it possible to differentiate between fixation-related artifacts and artifacts resulting from other preparation steps.

Figure 4 demonstrates the correlation of SPRM and EM for two cells. Figure 4a and Figure 4d show the results of the SPRM cell-substrate distance measurements *in vitro*. The dashed lines in Figures 4a, d indicate the cross sections that are used to evaluate the distance profiles shown in Figures 4b, e, respectively. Figures 4c, f show the SEM images along the same cross-sections as in Figures 4a, d (the contrasts in the SEM images have been inverted). The SEM crossections reveal several bright structures at the cell-substrate interface. Since the gray values within these structures are very similar to the gray values of the resin surrounding the cell, it can be concluded that they correspond to resin-filled cavities. As we will discuss later, the cell shown in Figures 4 d-f underwent the smallest changes in the cell-substrate distance during the chemical fixation (see Figure 5, cell 8). As this result suggests that the morphology of this fixed cell might be close to its physiological state, we chose to image this particular cell using STEM. For this purpose, we prepared a lamella from the center of the cell corresponding to the cross section shown in Figure 4d. As we can see in Figure 4g-j, the STEM images allow resolution of the cell membrane. Here, the resin appears bright and the cell membrane and the substrate relatively dark.

As we can see in the images recorded after the first sample mounting (Figure 4g, h), the resin was well attached to the substrate. However, after several mountings in the STEM, the fragile lamella disintegrated. Luckily, the resin detached cleanly from the gold surface of the substrate so that we can still investigate the cell-substrate interface. This is why, the surrounding of the resin appears very bright in Figures 4i, j.

Figure 4g shows an area with large adhering areas and small cell-substrate distances. As the separation between the intracellular and the extracellular space seems distinct, we conclude that the membrane is intact. Figure 4h instead shows a large oval structure close to the interface which resembles a vacuole. Here, we cannot tell with certainty whether this structure is closed or the membrane is torn. Therefore, the vacuole could already have been present in the living cell and have been preserved by fixation or introduced by further preparation steps.

Figures 4i, j show close-ups of two other cavities. While Figure 4i shows a distinct separation of the extracellular and intracellular space by the cell membrane, the edge of the cytosol in Figure 4j appears fuzzy suggesting that the membrane ruptured during the fixation or following preparation steps.

To quantitatively analyze the effect of chemical fixation of multiple cells, we determine the changes in SPRM measurements as described in the Experimental Section. The changes for all scanning points are then pooled for the individual cells and their probability density function estimated (see Figure 5). With the exception of cell 2, the median of the change in cell-substrate distance scatters around 0 *nm* within an interval of 10 *nm* without preferential directionality. This shows that the average distance between cell and substrate is maintained for the vast majority of measured cells. As for the changes in cytosolic RI, there is a clear trend towards higher values after fixation, which will be discussed below.

## 3 Discussion

In order to understand how glutaraldehyde mediated fixation can alter the cell-substrate interface, we will first discuss cell-substrate adhesion under physiological conditions. Then we will address the chemical processes during the fixation processes and how they might introduce changes at the cell-substrate interface.

The cells were seeded on a PLL-coated substrate. As PLL is positively charged under physiological conditions, negatively charged acidic side chains such as aspartic acid or glutamic acid in the membrane proteins can bind electrostatically [61]. The charged proteins in the membrane and cytosol are hydrated due to the polarity of the water molecules in the cytosol and within the culture media. Glutaraldehyde, which is often used in fixation protocols, links amino groups of close by polypeptides [61]. This causes crosslinking of membrane proteins to the PLL coating, however intracellular proteins inside the organelles or the cytoskeleton can also be crosslinked. During the crosslinking, the glutaraldehyde binds covalently to the amino acids of two closely spaced polypeptides. Consequently, amino groups lose their charges, forming a neutral complex with their two binding partners.

In our understanding, the change in charge distribution within the cell proteins might affect the tertiary structure of the proteins. The absence of the charges responsible for the hydration might result in a locally reduced water concentration.

The covalent bonds caused by crosslinking are much stronger than the electrostatic interaction responsible for the cell-coating adhesion in the living cell. Assuming that the system evolves in favor of energetically lower states during the incubation time of the crosslinker, we would expect changes in the cell-adhesion complexes when compared to the living cell. The change in the tertiary structure and the hydration of the proteins can also explain local changes in the cell-substrate distance. The observed increase of the cytosolic RI gives evidence for the validity of this explanation: As the RI of water (1.33) [62] is low compared to the RI of proteins (> 1.50) [63], a local reduction of water concentration would result in a higher cytosolic RI after the fixation. This explains the fixation-related contrast enhancement that we observed in the live images. Since changes in cytosolic RI lead to a shift of the entire reflection curve, the plasmon resonance angle shifts towards higher angles in areas of cell adhesion, while in uncovered areas the plasmon resonance angle remains constant.

The lateral resolution in SPRM is defined by the spot size of the illuminated area during scanning, which is diffraction limited (*d* ≈ 200 *nm*)[58, 49, 59]. The values measured with SPRM thus correspond to the average cell-substrate distance within the illuminated areas. In wide cavities it is therefore to be expected that EM images may show larger maximum values compared to SPRM measurements, while cavities narrower than its lateral resolution cannot be resolved. This explains why some of the cavities appear in the EM images and not in the SPRM results. However, in STEM we found a cavity wider than the lateral resolution of SPRM that does not appear in the corresponding measurements (see Figure 4j). The extensive preparation protocol carried out after fixation comprises 12 changes of the extracellular fluids (7x with EtOH, 5x with resin, both in increasing concentration) which result in diffusion processes across the cell membrane and therefore consecutive replacements of the intracellular fluids. This large number of diffusion processes is likely to change the morphology of the cell membrane and thus the shape and size of the cavities. In particular, Figure 4j shows a partially ruptured membrane which indicates that the diffusion across the cell membrane may not have worked ideally and therefore, this cavity can be considered an artifact.

Both methods have their obvious advantages and disadvantages: FIB-STEM offers unique resolution in x- and z-direction and also resolves cell organelles, whereas SPRM can display the situation in the living cell. One limitation of SPRM is its specificity for sample structure and optical properties of materials within the sample. The exchange of water with resin both in the cell-substrate cleft and within the cell leads to a strong increase in the RI of the samples, making it impossible to investigate morphological changes caused by this preparation step. Furthermore, after exposing the cells to *OsO*_4_ and uranyl acetate, the samples can only be handled in designated work areas and therefore not be further investigated with SPRM. However, the combination of these two methods allows us to benefit from all the above advantages in the same study.

## 4 Conclusion

We have shown a correlative workflow that enables us to complement FIB-S(T)EM studies with SPRM as an independent technique. This method has been successfully applied to a preparation protocol for one cell type and can serve to test and verify various fixation and preparation protocols in the future.

The comparison of the pre and post chemical fixation profiles of the adherent cell membrane allows us to exclude those cells that show a significant shift in the cell-substrate distance from subsequent EM investigations. This increases the likelihood that the cells studied in FIB-STEM are preserved in their natural shape. Our study shows that the average cell-substrate distance of the vast majority of investigated cells shows a shift of less than 10 *nm*. This suggests that the chemical fixation required for electron microscopy could maintain the structure at the cell-substrate interface which dispels one of the main objections to electron microscopy.

## 5 Experimental Section

### 5.1 SPRM

In the live-imaging mode, the sample is illuminated through the objective under the surface plasmon resonance angle using a projector as a light source. As a result, areas with close cell-substrate distance appear brighter than areas with higher cell-substrate distances or the surrounding media (see Figures 1b, 6a).

In scanning SPRM, we focus radially polarized laser light at the glass-gold interface which results in an illumination of the sample with p-polarized light under a broad range of angles. Most of the incident light is reflected at the interface and is captured by the objective whereas other parts of the incident light are absorbed and excite surface plasmons. By capturing images of the back focal plane (BFP) of the objective (see Figure 1e), we can extract the angle spectrum of the reflected light (see Figure 1f). This data can be analyzed based on the transfer-matrix method. Here, the sample is modelled as a multilayer system with specific optical properties (see Figure 1d). The metal layers as well as the cell membrane are assumed constant whereas the thickness of the culture media layer as well as the intracellular RI are variable. These two variables are obtained by fitting the simulated curve to the data (see Figure 1f) [56]. As a result of this analysis, a cell-substrate distance profile as well as the profile of the cytosolic RI are generated for each scan.

### 5.2 *Align and compare* software

The software *align and compare* was written in MATLAB and allows us to determine the fixation induced changes in the cell-substrate distance and in the cytosolic RI. Before starting the *align and compare*-routine, we analyzed the scanning data as briefly described before.

This analysis is based on the description of the sample as a multilayer system. The reflectance of such a system can be described as a function of the angle of incidence using the transfer-matrix method [64]. As we showed before, this method can be used to analyze the intracellular RI separately from the cell-substrate distance by optimizing the overlay of the simulated and the measured reflectance [56]. As a result of this analysis, the cell-substrate distance as well as the RI are saved as individual data sets.

After this initial analysis, the cell-substrate distance profiles and RI profiles which correspond to the measurements before and after the fixation of the same cell are loaded into the *align and compare* routine. The software displays the distance profiles corresponding to the two states (see Figure 6b, c). We now identify three salient points in the two distance profiles via graphical input. The coordinates of these inputs build the basis for the coordinate transform which enables us to transform the scanning data of the two measurements into a common system. The coordinate transformation consists of a shift and a rotation in the x-y plane. By applying this transformation, we transfer the data corresponding to the measurement after the fixation into the coordinate system of the first measurement (see Figure 6d). Afterwards, the user is asked to mark the cell-adherence area in the scan based on graphical input. A SPRM live-image (see Figure 6a) helps to identify the respective areas. Now, a polygon including all the points entered by the user (see Figure 6e) is determined and the areas outside the polygon are discarded. Finally, the changes in the distance and RI inside the polygon are calculated point wise. These results are plotted in form of area plots (see Figure 6f), but also as line plots between two points chosen by the user via graphical input (see Figure 3 and Figures 4 a, b, d, e) as well as in form of violin plots (see Figure 5) using Python.

### 5.3 Fixation and preparation for FIB

The fixation, heavy metal stainings and resin embeddings have been carried out based on the protocol established by Belu *et al.* [7]. Unless stated otherwise, all the chemicals have been purchased from Sigma-Aldrich.

#### 5.3.1 Chemical fixation

First, the samples were rinsed three times with 37 °C warm PBS. The cells were fixed using 37 °C warm 3.2% glutaraldehyde solution at room temperature (RT) for 15 min. Afterwards, the samples were washed three times with PBS at RT.

#### 5.3.2 Staining with osmium tetroxide and uranyl acetate

The samples were rinsed twice with MilliQ at RT. Afterwards, the samples were incubated with 1% OsO_4_ in cacodylate buffer (Morphisto) for 2 *h* on ice. The samples were rinsed 5x with MilliQ before they were exposed to 1% tannic acid (Electron Microscopy Science) for 30 *min* at RT and again rinsed five times with MilliQ. Afterwards, the samples were incubated with 2% uranyl acetate at 4°C overnight and rinsed again five times with MilliQ.

#### 5.3.3 Ethanol Dehydration

The cells were rinsed with gradual increasing ethanol (EtOH) series at RT: 5 *min* each with 10% EtOH, 30% EtOH, 50% EtOH. 15 *min* with 70% EtOH, three times 5 *min* each with 90% EtOH and 95% EtOH and finally 5 *min* with 100% EtOH.

#### 5.3.4 Resin embedding

The resin used for this embedding was prepared as follows: 20 *ml* DDSA were mixed with 12.5 *ml* of EPON 812 and 17.3 *ml* MNA were mixed with 15.2 *ml* EPON 812. Afterwards, the two solutions were mixed. 1.3 *ml* of DMP30 were added to 32.5 *ml* of this mixture and stirred for 1 *h*.

The samples were exposed to resin-EtOH solutions with gradual increasing resin fraction at RT: We started the resin embedding by incubating the sample to a resin solution with 3:1 ratio (EtOH:resin) for 3 *h*, followed by 3 *h* incubation with a 2:1 solution, an overnight incubation with a 1:1 solution, a 3 *h* incubation with a 1:2 solution, a 3 *h* incubation with a 1:3 solution and a 3 *h* incubation with pure resin. Afterwards, EtOH was splashed repeatedly over the sample from the sides of the dish to remove the supernatant. Finally, the resin was cured at 60°C for 12 *h*.

### 5.4 Metal deposition

Before mounting the samples in the FIB-SEM system, we sputtered a platinum layer on the sample sample surface to avoid charging effects (using Balzer SCD004, 30 s, 30 mA).

Using a FIB-SEM system (Helios NanoLab 600i, FEI Co., Hillsboro, OR, USA), we localized the cells and deposited an additional thin platinum layer on the region of interest via electron beam induced deposition to protect the sample from damages caused by the ion source. Before starting the FIB sectioning, we deposited an additional platinum layer of several micrometer thickness which stabilized the sample during the FIB sectioning using ion beam induced deposition.

### 5.5 EM

The SEM imaging as well as the FIB sectioning were conducted on a Helios NanoLab 600i (FEI Co., Hillsboro, OR, USA). For STEM, a thin lamella (thickness at the cell-substrate interface ≈100 *nm*) was cut from the center of the cell and soldered to a TEM grid. The STEM Imaging of the lamella was conducted on a Magellan 400 FESEM (FEI Co., Hillsboro, OR, USA) with electron beam energy of 28keV and current of 0.1 nA. For high-contrast, high-resolution micrographs a STEM II detector in bright field (BF) mode was used.

### 5.6 SPRM sample preparation

The samples consist of a high index coverslip which is coated with a 3 *nm* thick chromium layer and a 38−40 *nm* thick gold layer. Specific markers which allow for a unique identification of each position on the substrate are fabricated on top of the gold layer using photo resist. These substrates are glued to 35 *mm* polystyrene Petri dishes with a hole in the bottom and can afterwards be used as substrates for cell culture.

#### 5.6.1 Substrate cleaning

Used substrates: High index coverslips (Olympus, HIGH INDEX-CG)

Clean substrates with ethanol, aceton and isopropanol (5 *min* each).

O2-plasma cleaning

machine: Barrel Reactor TePla Gigabatch 310M

parameters: 200 *W*, 50 *sccm*, 5 *min*

#### 5.6.2 Chromium deposition

3 *nm* Cr sputtered

machine: Leybold Univex 450C

parameter: 300 *W* DC, 60 *sccm* Argon (2.1 · 10^−2^ *mBar*), 5.3 *s*

#### 5.6.3 Gold deposition

Gold evaporation directly after the chromium deposition

machine: PLS-570

parameters: 38nm Au

#### 5.6.4 Lithography

Baking of samples 5 *min*, 130°C

Spincoating: AZ5214E: 4000 *rpm*

Baking of samples 1 *min*, 110°C

Mask Aligner MA4: 2.3 *s*, 75 *mW/cm*^2^, hard contact

machine: Süss MA/BA 8

Development: MIF326, 45 *s*, stop in DI-water ∼5 *min* until guide value > 10 *MOhmcm*

Baking of samples 10 *min*, 135°C

### 5.7 Neuronal cell culture

200 000 primary neurons were suspended in 2.5 *mL* and seeded on the PLL coated substrates The entire media was replaced 3h after plating.

Half the volume of the media was replaced each 3 or 4 days (alternating).

#### 5.7.1 Culture media

10 *mL* Neurobasal Medium (from Life Technologies, 21103049)

100 *µL* B-27 supplement (from Life Technologies, 17504044)

25 *µL* L-Glutamine (from Life Technologies, 25030024)

10 *µL* Gentamicin, (from Sigma, G1397)

## Contributions

E.K. conceived, executed and evaluated the SPRM experiments, wrote the software *align and compare* and wrote the manuscript. S.S. conceived and executed the heavy metal stainings and the resin embedding. H.H. wrote the SPRM analysis software. E.B-R and E.N. conceived and executed the FIB-S(T)EM experiments. A.O. supervised the project. All authors reviewed the manuscript.

## Acknowledgement

The authors thank Klaus Kreysing, Dr. Kristian Franze and Dr. Moritz Kreysing for fruitful scientific discussions, Michael Prömpers for the sample production and Bettina Breuer and Timm Hondrich for the neuron isolation.

